# Oligomycins Inhibit *Magnaporthe oryzae Triticum* and suppress wheat blast disease

**DOI:** 10.1101/2020.05.13.094151

**Authors:** Moutoshi Chakraborty, Nur Uddin Mahmud, Abu Naim Md. Muzahid, S. M. Fajle Rabby, Tofazzal Islam

## Abstract

Oligomycins are macrolide antibiotics, produced by *Streptomyces* spp., show biological activities to several microorganisms like bacteria, fungi, nematodes and peronosporomycetes. Conidiogenesis, germination of conidia and formation of appressoria are crucial for a successful disease cycle and pathogenicity of the filamentous fungal phytopathogen. The goal of this research was to evaluate the effects of two oligomycins, oligomycin B and oligomycin F along with a commercial fungicide Nativo^®^ 75WG on hyphal growth, conidiogenesis, conidia germination, appressoria formation, and disease development of a worrisome wheat blast fungus, *Magnaporthe oryzae Triticum (MoT)* pathotype. Both oligomycins suppressed the growth of *MoT* mycelia depending on the dose. Between the two natural products, oligomycin F displayed the maximum inhibition of *MoT* hyphal growth accompanied by oligomycin B with minimum inhibitory concentrations (MICs) of 0.005 and 0.05 μg/disk, respectively. The application of the compounds also completely halted the conidia formation in the *MoT* mycelia in agar medium. A further bioassay showed that these compounds significantly inhibited *MoT* conidia germination and induced lysis; if germinated, induced abnormal germ tube and suppressed appressoria formation. Interestingly, the application of these macrolides significantly inhibited wheat blast disease on detached leaves of wheat. This is a first report on the inhibition of mycelial growth, process of conidia formation, germination of conidia, morphological changes in germinated conidia, and suppression of blast disease of wheat by oligomycins from *Streptomyces* spp. A further study is needed to evaluate the mode of action and field trials of these natural compounds to consider them as biopesticides for controlling this devastating wheat killer.

## Introduction

Oligomycins are macrolide antibiotics, produced by some strains of *Streptomyces* species. They have a broad-spectrum of biological activities to organisms like fungi, bacteria, nematodes and peronosporomycetes [1–4]. The *Streptomyces* spp. are common soil-dwelling bacteria, which have been broadly used as bio-control agents [5]. *Streptomyces* species produce a number of bioactive compounds, such as antifungal, antiviral, antibacterial, anticancer, nematicidal, and antioxidant properties [5, 6]. Several previous studies show that the effectiveness of some strains *Streptomyces* spp. in biological control of phytopathogens is largely related to the production of oligomycins [7]. The oligomycins are mitochondrial F1F0 ATP synthase inhibitors that cause apoptosis in a number of cell types [8]. The oligomycin complex which was first documented in 1954 in a strain of a soil bacterium, *S. diastatochromogenes* was highly inhibitory against fungi [1]. Antifungal, antitumor, insecticidal, immunosuppressive and nematocidal properties of oligomycins have been reported [1–3, 7, 9]. The oligomycins contain the analogs / isomers A through G. Different isomers are highly selective for disrupting mitochondrial metabolism [3, 4, 8, 10]. Although the biological activities of oligomycins on fungi and peronosporomycetes have been reported, nothing is known about the effect of these natural products on a notorious wheat blast fungus *Magnaporthe oryzae Triticum* (*MoT*). The bioactivities of oligomycins to different classes of fungal species indicates that their targets may involve varieties of cellular processes, such as inhibition of mycelial growth of *Cladosporium cucumerinum, Magnaporthe grisea, Colletotrichum lagenarium, Botrytis cinerea, Cylindrocarpon destructans, Fusarium culmorum, Erysiphe graminis* and *Phytophthora capsici* [3, 11], lysis and motility inhibition of downy mildew zoospores (*Plasmopara viticola*) of grapevine, and inhibition of motility of zoospores of damping-off phytopathogen, *Aphanomyces cochlioides* [4].

The wheat blast fungus *MoT* is one of the most damaging pathogens of wheat [12–15]. The fungal three-celled, hyalin, pyriform conidium is bound to the host surface by an adhesive secreted from the tip of the conidium [14, 16, 17]. The conidium attached grow a germination to form a hyphal germ tube, an aspersorium and a penetration peg, which completes the infection by rupturing the cuticle, allowing it to penetrate the epidermis of the host [16, 18]. The invasion of plant tissue happens by bulbous hyphae invaginating the host plasma membrane and invading the epidermal cells [16–18]. The fungus can attack wheat plants at any stages of development and infects the aerial parts of wheat plants, including leaves, nodes, stems, and spikelets [15, 17, 19]. Mycelium can survive in embryo, endosperm, and kernal tissues of wheat seed. Wheat blast mainly affects wheat heads; it bleaches the infected heads, resulting deformed seed or no seed production [14]. The badly affected wheat head can die, leading to a drastic reduction in grain yield. Bleaching of spikelets and entire head at an early stage is perhaps the most common recognizable symptom [12, 14, 15]. Infected seeds and airborne conidia usually transmit the disease, and the fungus may survive in infected seeds and crop residues [20]. Therefore, pyriform conidia developed from conidiophores and conidia germination with appressorial development at the germ tube tips are essential steps of the disease cycle of *MoT* [16]. Disrupting any of these asexual life stages reduces the chance of pathogenesis [21]. Finding of natural bioactive compounds inhibiting any of these phases of asexual life is considered to be the first step in the production of a new fungicide for *MoT*.

Wheat blast was first recorded in Brazil in 1985, and has been important locally since its introduction in Brazil, Bolivia and Paraguay [12, 19]. The cultivation of wheat (*Triticum aestivum*) was increased in Bangladesh, making it the 2^nd^ largest food source after rice. In 2016, a sudden outbreak of wheat blast disease occurred in Bangladesh which was the first incident in any countries outside of South America [14, 22]. About 15,000 hectares of wheat crops have been destroyed, resulting in about 15% crop losses in Bangladesh [14]. The outbreak of wheat blast concerns crop scientists as it extends further to major wheat-producing regions in neighboring South Asian countries and Africa due to similar climatic conditions in such regions [23]. Plant pathologists have cautioned that this disease is expected to transmit to India, Pakistan and China, that are ranking 2^nd^, 8^th^ and 1^st^ in the world wheat production, respectively [15, 24].

Current wheat blast disease management methods include the utilization of synthetic fungicides. Most of the synthetic fungicides are harmful to the environment and health of living species including humans [25–26]. Indiscriminate application of synthetic commercial fungicides in plant protection results in resistance to fungicides [13, 17]. In Brazil and other South American countries, some *MoT* strains showed resistance to these chemical arsenals due to the extensive use of strobilurin (QoI) and triazole fungicides [27, 28]. Nowadays, natural products that are environment-friendly and have minute toxicity to living organisms are gaining popularity as important sources for producing ecologically suited alternative fungicides for protecting plants. Therefore, search for novel bioactive natural products against *MoT* is needed.

The biological approach for the plant disease management offers a better alternative to the control of wheat blast disease owing to its safety for human use and the environment. So far, no report has been demonstrated on the antagonistic impacts of *Streptomyces* spp., and/or their secondary metabolites to control wheat blast disease. We screened 150 natural compounds belonging to the class alkaloids, terpenoids, macrolides, macrotetrolides, tepenoids, and phenolics isolated from different plants and microorganisms on mycelial growth and asexual development of wheat blast fungus *MoT* in our laboratory [21]. Among them, two macrolides, oligomycin B and oligomycin F, previously extracted from the marine *Streptomyces* spp. [3–4] displayed very strong inhibitory effects against *MoT* both *in vitro* and *in vivo*. This is known to be the first report of the bio-control of the notorious wheat blast disease by some macrolides from the *Streptomyces* spp. The specific objectives of this study were to: (i) test the effects of oligomycins B and F on mycelial growth of *MoT*; (ii) examine the effects of these macrolides on conidiogenesis, conidia germination and their subsequent morphological transitions in sterilized water medium; and (iii) evaluate the effects of these macrolides on the suppression of the development of wheat blast disease in the detached wheat leaves.

## Materials and methods

### Chemicals

Oligomycin B and oligomycin F (Fig. 1) were isolated from the marine bacteria, *Streptomyces* sp. strains B8496, B8739 and A171 [3–4]. These compounds were generously provided by Dr. Hartmut Laatsch of Georg-August University Goettingen, Germany. Fungicide Nativo^®^ WG 75 was purchased from Bayer Crop Science Ltd.

**Fig 1.**
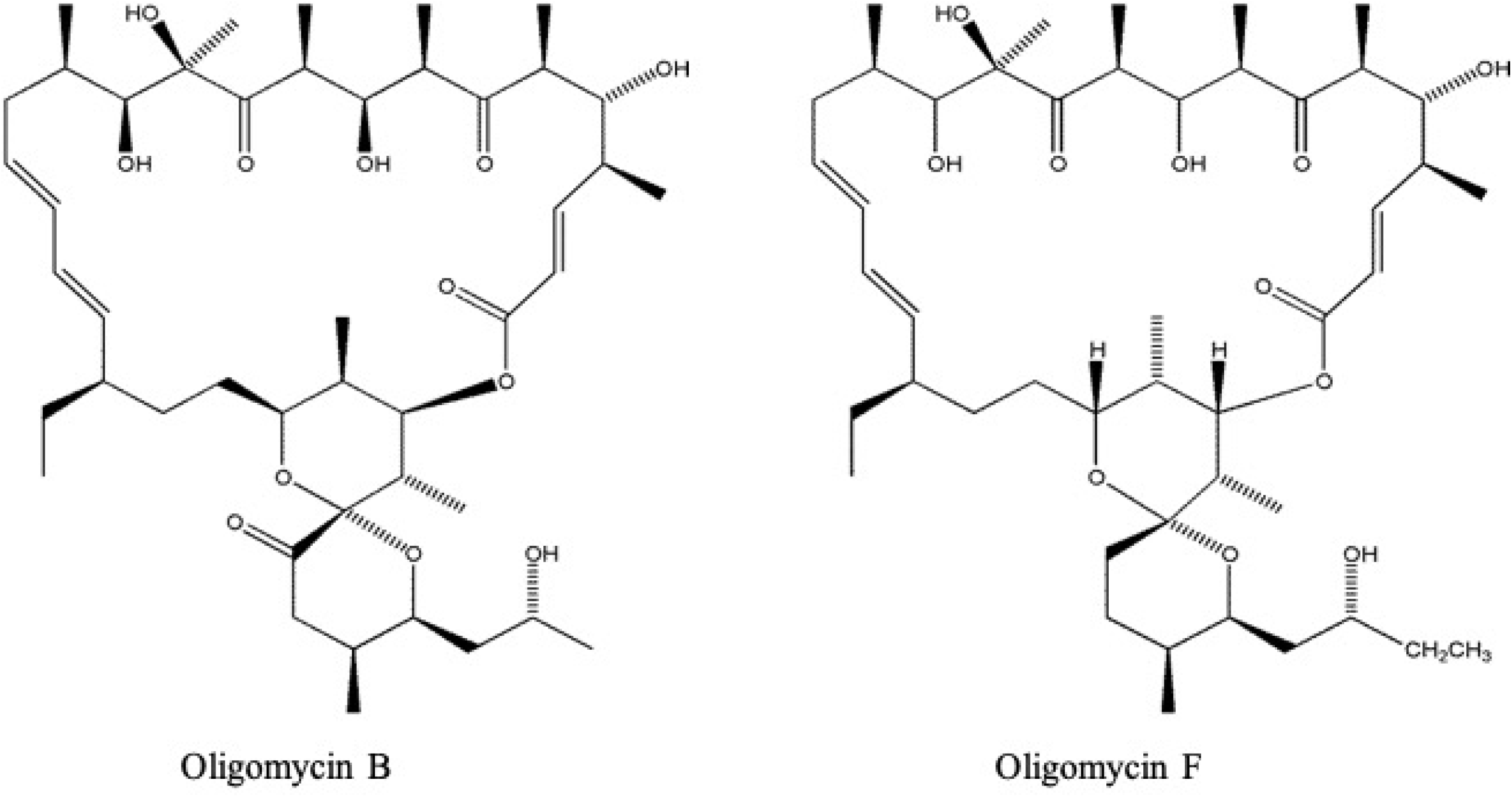
Structure of macrolides isolated from *Streptomyces* spp. tested toward *M. oryzae Triticum (MoT)* (Dame et al., 2016)

### Fungal strain, media and plant materials

The strain BTJP 4 (5) of *MoT* was isolated from the spikelets of wheat cv. BARI Gom-24 (Prodip) with blast infection, which was collected from a highly infected field in Jhenaidah, Bangladesh in 2016. The isolated strain was conserved at 4°C on dry filter paper before used in the present research (Islam et al., 2016). It was re-cultured on the PDA (Potato Dextrose Agar 42 g/L) medium and incubated at 25°C for 7-8 days. After placing a small block of *M. Oryzae Triticum (MoT)* pathotype isolate BTJP 4 (5) in the above-mentioned medium, the plate was incubated at 25°C [14].

For sporulation, 10-days-old fungal cultures grown on PDA medium were washed with 500 ml of deionized distilled water for removing aerial mycelia and then kept at room temperature (25-30°C) for 2-3 days [14, 29]. The conidial and mycelial suspensions were filtered by two layers of cheese cloth and adjusted to a concentration of 1 × 10^5^ conidia/ml. Conidia were collected and put in water. Conidial germination was visualized under a microscope and germinated conidia numbers were counted. BARI Gom-24 (Prodip) wheat cultivar which is susceptible to *MoT,* was used for the disease suppression bioassay. Wheat leaves were separated from the seedlings of five-leaf stage for bioassay of *in vivo* disease suppression [30].

### Mycelial growth inhibition by oligomycins

The *in vitro* antifungal efficacy of pure oligomycins and commercial fungicide, Nativo ^®^ WG 75, were evaluated on the basis of the hyphal growth inhibition rate of *MoT* isolate BTJP 4 (5) by the modified Kirby - Bauer disk diffusion technique [31]. A series of concentrations of both the natural compounds were prepared in ethyl acetate. However, Nativo ^®^ WG75 was added in distilled water to make fungicidal solution. Nine-millimeter diameter of filter-paper disks (Sigma-Aldrich Co., St. Louis, MO, USA) were filled with bioactive compound suspensions at concentrations ranging from 0.005 to 2 μg / disk. Paper disks were placed on one side of the Petri dish (2 cm from the edge) comprising of 20 ml of PDA. Five millimeter diameter mycelial plugs from seven-days-old PDA cultures of *MoT* were mounted on the opposite side of Petri dishes (9 cm diameter) perpendicular to paper disk. As a positive control, Petri dishes inoculated with fungal mycelia plugs against fungicide Nativo ^®^ WG75 were used, while fungal colonies were used as a negative control without any treatment. The minimal inhibitory concentration (MIC) of oligomycin B and F was also measured. After 10 days, the antimicrobial activity was shown by the diameter of growth inhibition zones. For each concentration used, the experiment was replicated five times. Plates were incubated at 25°C until fungal mycelia fully covered the agar surface of control plate. The fungal colony’s radial growth was measured in centimeter (cm) with a meter ruler along with two diagonal lines drawn on each plate’s opposite side. Data were recorded by the measurement of pathogen inhibition zone and mycelial growth, and the percentage of inhibition of radial growth (PIRG) (± standard error) [32] was calculated from mean values as:

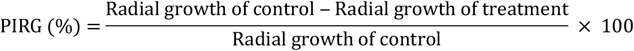

Hyphal morphologies in the vicinity of compounds were observed under a ZEISS Primo Star microscope at 40x and 100x magnification. Images of inhibition zone were captured through a digital camera (Canon DOS 700D) and images of morphologies of hyphae were recorded by a microscopic camera (ZEISS Axiocam ERc 5s).

### Conidiogenesis inhibition by oligomycins

The stock solution of each oligomycin was prepared in 10 μl of dimethyl sulfoxide (DMSO). Each compound solution was then made ready in distilled water at concentrations of 5, 10 and 100 μg / ml. The final concentration of DMSO was never higher than 1% (v / v) for the working solution, which does not affect the hyphal growth or sporulation of *MoT*. Preparation of 5 ml fungicidal suspension of Nativo^®^WG75 was carried out in distilled water (5, 10 and 100 μg/ml) for using as a positive control. For conidia formation (i.e., conidiogenesis) observation, mycelia were washed out from the Petri dish of 10-days-old PDA culture of *MoT* for reducing nutrient to induce conidiogenesis in the fungal mycelia [14, 29]. Exactly 50 μl of each compound and Nativo^®^WG75 were exerted on the 10 mm *MoT* mycelial agar block at 5, 10 and 100 μg/ml put in Nunc multidish (Nunc). Only sterilized water was exerted on the *MoT* fungal mycelial agar block with 1% DMSO which used as negative control. Dishes with mycelial agar blocks of *MoT* fungus were incubated at 28°C with optimum humidity and lights. Formation of conidia was visualized under a ZEISS Primo Star microscope at 40x magnification after 24 h, and the images were captured with a microscopic camera (ZEISS Axiocam ERc 5s). The experiment was replicated five times, and five replications were performed each time.

### Conidial germination inhibition and morphological changes of germinated conidia by oligomycins

Preparation of stock solution of each oligomycin (0.1 μg) was done in 10 μl of dimethyl sulfoxide (DMSO). Exactly 0.1 μg/ml concentration of each compound was then achieved by adding distilled water. Nativo^®^WG75 solution was prepared with distilled water at 0.1 μg/ml which served as a positive control. The conidia germination bioassays were carried out by following the protocol of Islam and von Tiedemann [33]. Briefly, 100 μl from 0.1 μg/ml sample solution was added directly to 100 μl conidial solution (1 × 10^5^ conidia) of *MoT* to make a final volume of 200 μl (0.05 μg/ml sample solution) which poured into a well of a plant tissue culture multi-well plate. Afterwards, the solution was quickly mixed with a glass rod and incubated at 25°C. The presence of ≤1% DMSO in the conidial suspension was maintained to avoid any effect on conidial germination or further developments of the germinated conidia. Sterilized water was used as control with 1% DMSO. The multidish containing conidial sample solution in wells was incubated in a moisture chamber at 25°C for 6 h, 12 h and 24 h without the provision of light. A total of 100 conidia from each of five replicates were examined under bright field microscope (ZEISS Primo Star, Germany) at 100x magnification to evaluate the percentage of conidial germination and major developmental transitions of the treated conidia, and the pictures were captured with a microscopic camera (ZEISS Axiocam ERc 5s). The experiment was performed five times and each time it was repeated in five replications. In time wise microscopic observation, the treatments of conidia with tested compounds likely result in germination, no germination, alteration of the morphology of germ tubes, appressoria and/or mycelial growth. The percentage of conidia germination (± standard error) was calculated from mean values as: CG % = (C - T)/C × 100; Where, CG = conidial germination, C = percentage of germinated conidia in control, and T = percentage of germinated conidia in sample. Same calculation was used for other conditions mentioned.

### Suppression of wheat blast disease on detached leaves by oligomycins

As stated earlier, stock solutions of oligomycins B and F were prepared using dimethyl sulfoxide (DMSO). Then preparation of 5, 10 and 100 μg/ml concentrations of each compound was carried out in distilled water where the final DMSO concentration never exceeded 1%. Different concentrations (5, 10 and 100 μg/ml) of Nativo^®^WG75 were served as positive control. Sterilized water was applied as negative control with 1% DMSO. From the five-leaf stage seedlings, wheat leaves were separated and introduced in plates lined with moist absorbent paper. Three 20-μL droplets of the formulated compounds were then inoculated on each leaf. Afterwards, the leaves were permitted to absorb the compounds for 15 minutes. Consequently, 1 × 10^5^ *MoT* conidia were introduced onto the leaf surface. With 100% relative humidity, each of the dishes was incubated at 28 °C, first 30 h in dark condition, then 2 days in continuous light condition. The test was performed five times independently. The wheat blast lesions diameter was measured from 12 leaves per experiment.

### Statistical analysis, experimental design/replications

Experiments were performed using a complete randomized design (CRD) to determine the biological activities of the pure compounds. Data were evaluated by a one-way variance analysis (ANOVA), and the mean values were distinguished by the posthoc statistic of Tukey’s HSD (honest significant difference). All the statistical analyses were carried out by using SPSS (IBM SPSS statistics 16, Georgia, USA) and Microsoft Office Excel 2010 program package. Mean value ± standard error of 5 replications were used in Tables and Figures.

## Results

### Oligomycins inhibit mycelial growth and induce alterations in hyphal morphology of *MoT*

150 secondary metabolites of several plants and micro-organisms have been evaluated to find out whether the natural products inhibit *MoT* fungal (Chakraborty et al. 2020). Among these natural compounds evaluated, two oligomycins (B and F) (Fig 1) extracted from the *Streptomyces* spp. showed significant hyphal growth inhibitory activities against *MoT* fungus in the PDA medium. Fig 2 demonstrates the inhibition of *MoT* hyphal growth by the tested oligomycins. Between these two compounds, oligomycin F depicted the strongest mycelial growth inhibitory activity against the isolate BTJP 4 (5) of *MoT.* The highest *MoT* hyphal growth inhibition was observed by oligomycin F followed by oligomycin B. Both compounds demonstrated potency on the tested phytopathogen ranging from 57.1 ± 1.3% to 73.9 ± 2.5% at 2 μg/disk (Fig 3). The activities of these macrolides were compared with that of the commercial fungicide used in the field, Nativo^®^ WG 75 whose control percentage was 81.9 ± 0.9% (at 2 μg/ disk).

**Fig 2.**
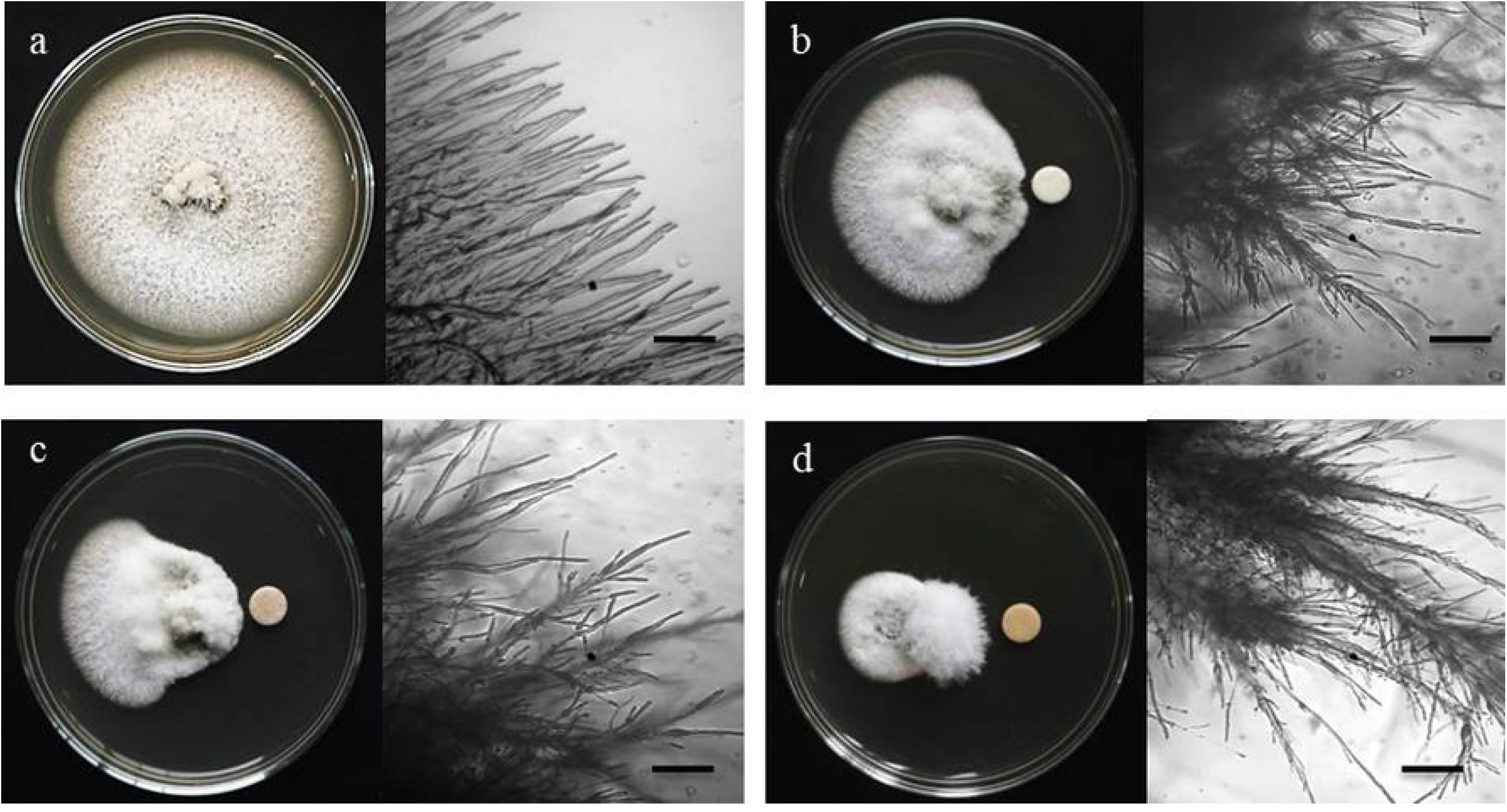
Macroscopic and microscopic view of *in vitro* antifungal activity of oligomycin B, oligomycin F and a commercial fungicide nativo^®^ WG75 against *M. oryzae Triticum (MoT)* at 2 μg/disk. (a) Control, (b) Oligomycin B, (c) Oligomycin F, (d) Nativo^®^ WG75. Bar = 50 μm.

**Fig 3.**
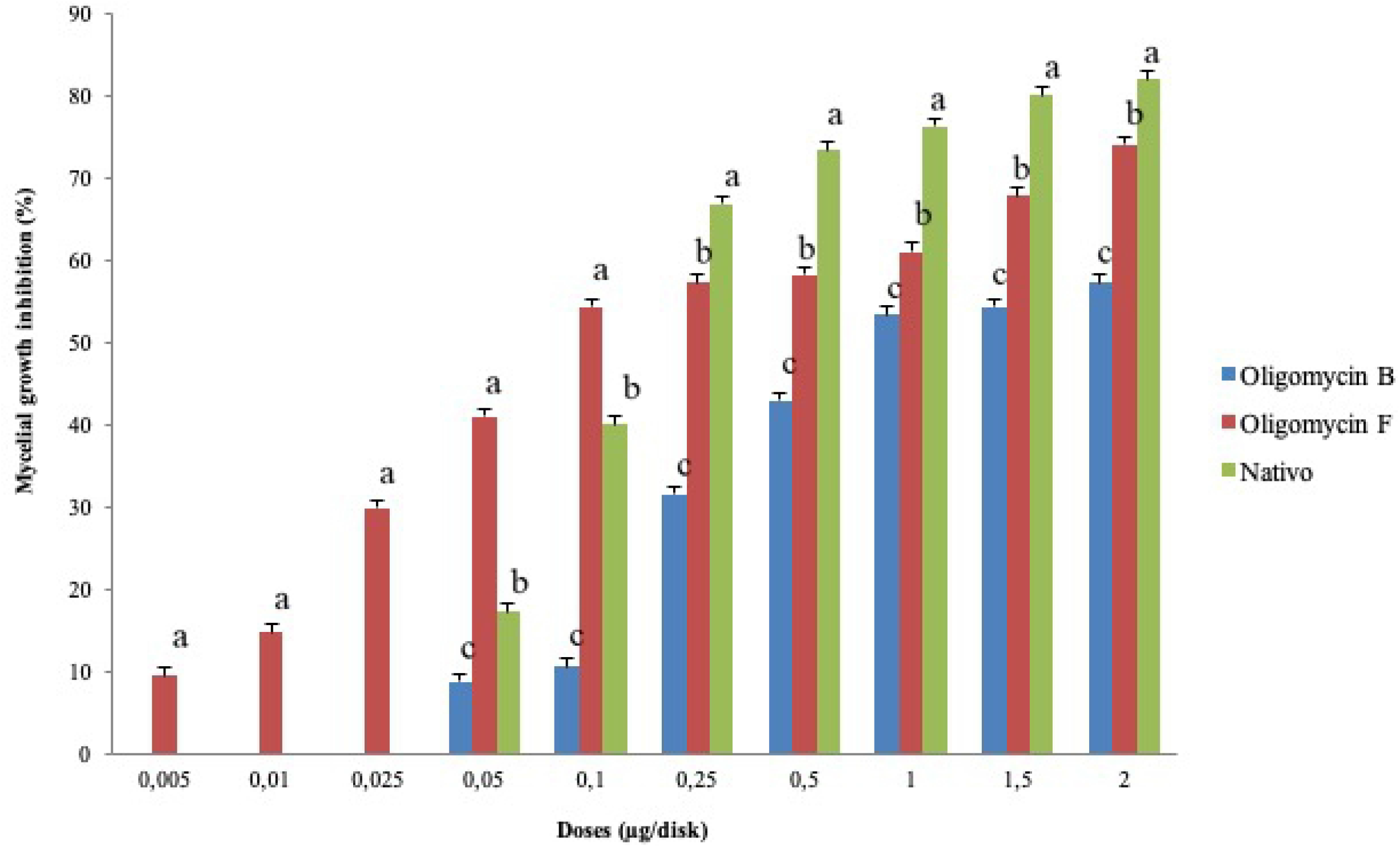
Inhibitory effects of oligomycin B, oligomycin F and a commercial fungicide nativo^®^ WG75 on hyphal growth of *M. oryzae Triticum (MoT)* in PDA media. The data are the mean ± standard errors of five replicates for each dose of the compound tested at a 5 % level based on the Tukey HSD (Honest Significance Difference) post-hoc statistic.

All the macrolides inhibited *MoTs* mycelial growth, though the strength varied. With the increase in oligomycin concentration the suppressive consequences increased from 0.005 to 2 μg/disk (Fig. 3). Neither of the macrolides, however, showed any activity against *MoT* at concentration lower than 0.005 μg/disk. Oligomycin F showed extensive inhibition of the pathogen at 2 μg/disk (73.9 ± 2.5 %) followed by 1.5 μg/disk (67.6 ± 0.9 %) and 1 μg/disk (60.9 ± 2.5%). The suppression percentage of oligomycin B were 57.1 ± 1.3 %, 54.3 ± 1.3% and 53.3 ± 1.5%, at 2, 1.5 and 1 μg/disk, respectively. Minimum inhibitory concentrations (MICs) of oligomycin F and oligomycin B 0.005 and 0.05 μg/disk (Fig 4) were also able to inhibit hyphal growth at 11.4 ± 2.3% and 8.63 ± 1.3%, respectively. Compared to these macrolides, minimal inhibitory concentration of Nativo was 0.05 μg/disk (Fig 4). At varying concentrations, the mycelial growth inhibitory activities of these macrolides similar and sometime significantly stronger than fungicide Nativo^®^ WG 75. Oligomycin F displayed inhibitory activity against *MoT* about 10-fold lower concentration than those of Nativo^®^ WG 75 and oligomycin B.

**Fig 4.**
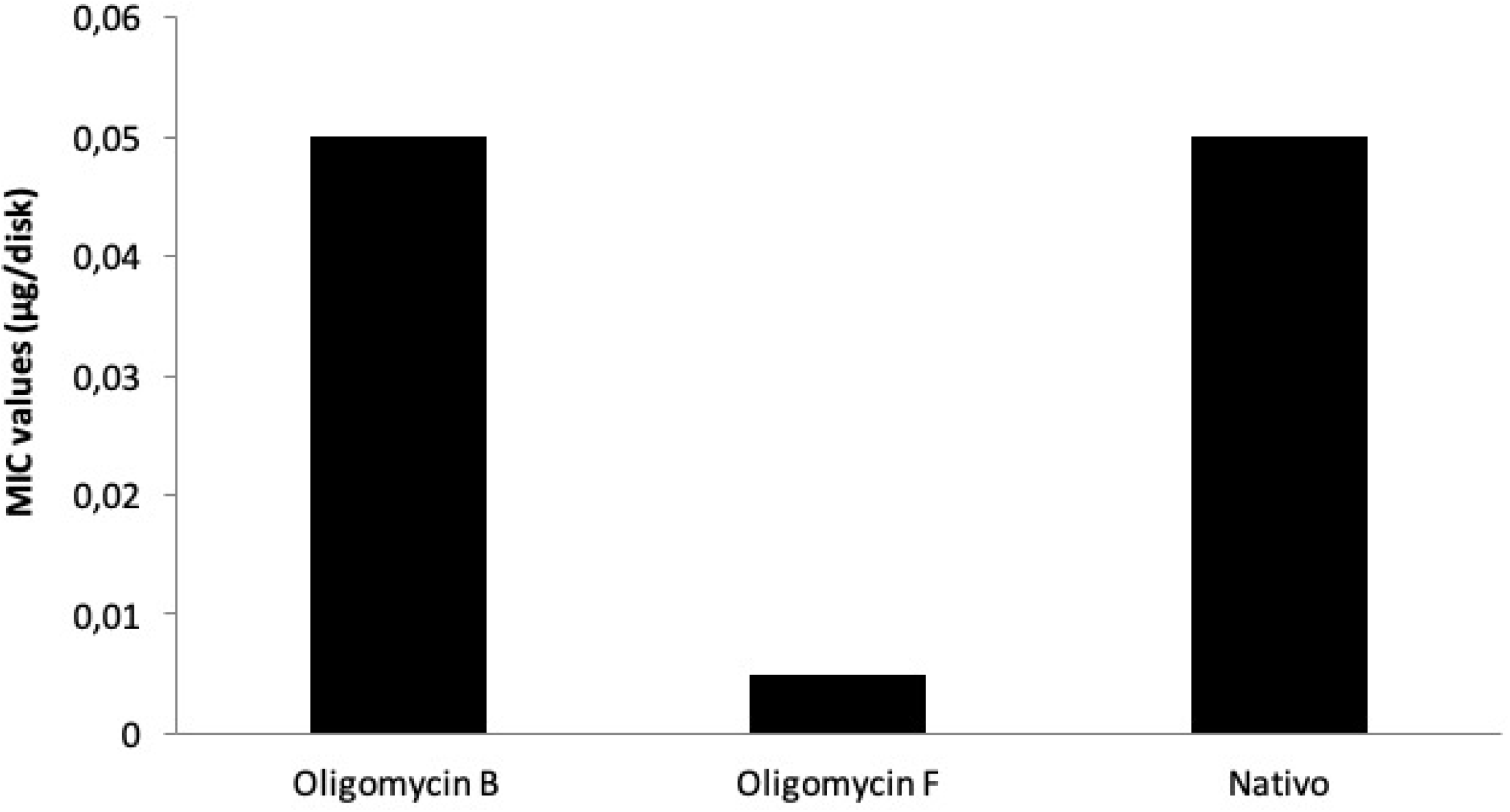
Minimal Inhibitory Concentration (MIC) values of oligomycin B, oligomycin F and a commercial fungicide nativo^®^ WG75 on hyphal growth of *M. oryzae Triticum (MoT)* in PDA medium.

Microscopic studies depicted that the untreated *MoT* hyphae had polar and tubular growth with smooth, branched, hyaline, plump, septate, and intact hyphae (Fig 2a). On the other hand, the morphology of hyphae treated with natural macrolides was apparently different from the normal hyphae of control. In contrast to the control group tubular hyphae, a noteworthy increase in branch frequency per unit length of hyphae, destruction of regular growth with twisted ridges and corrugations, and irregular swelling of hyphal cells were found when *MoT* was treated with oligomycins (Fig 2b and 2c). The *MoT* hyphae showed crystal appearance, loss of polar growth, erratic branching and swelling with twisted ridges and corrugations in the vicinity of Nativo ^®^ WG75 (Fig 2d). Morphological changes of *MoT* by two oligomycins were different from those occurred by the effect of commercial fungicide, Nativo^®^WG75

### Oligomycins suppressed conidiogenesis in *MoT*

We also tested whether the oligomycins (B and F) have an effect on the process of conidia production called conidiogenesis from the *MoT* mycelia. Conidiogenesis of *MoT* was inhibited by both the compounds. Huge conidia formation was found in the hyphal agar block of negative control after 24 h of incubation (Fig 5a). In the hyphal agar block of positive control, conidiogenesis inhibited in decreasing manner at 5, 10 and 100 μg/ml (Fig 5d). Compared to negative control, conidia formation was remarkably decreased by the oligomycins at 5 and 10 μg/ml concentrations (Fig 5b and 5c). Both the compounds and the positive control Nativo^®^WG75 completely blocked conidiogenesis at 100 μg/ml (Fig 5b and 5c). Moreover, microscopic findings demonstrated the existence of broken mycelial tips with no sign of conidiophore and conidial growth on the surface of the agar medium treated with macrolides at 100 μg / ml.

**Fig 5.**
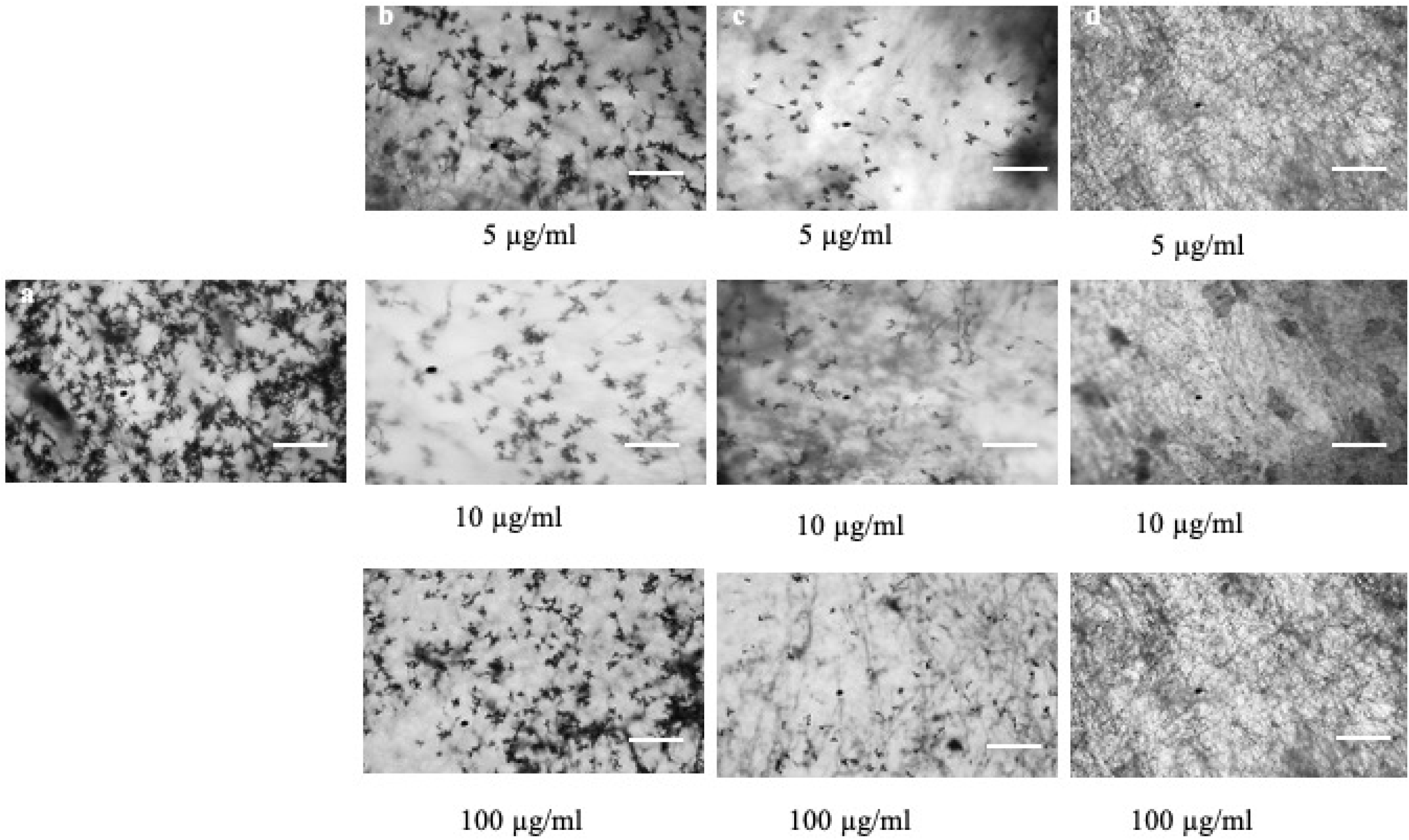
Effects of oligomycin B, oligomycin F and a commercial fungicide Nativo^®^ WG75 on the inhibition of conidiogenesis of *M. oryzae Triticum (MoT)* in Nunc multidish at 5 μg/ml, 10 μg/ml, 100 μg/ml. (a) Control, (b) Oligomycin B, (c) Oligomycin F, (d) Nativo^®^ WG75. Bar = 50 μm.

### Inhibitory effects of oligomycins on conidial germination of *MoT*

Oligomycin B and oligomycin F at 0.05 μg / ml were used to determine the inhibition of conidial germination of *MoT* in a multi-well plates. After 6 h of incubation percentage of conidial germination was recorded (Table 1). Conidial germination was 100% in water, but conidial germination was 50.3±0.7% in the case of Nativo ^®^ WG75. Both macrolides, oligomycin B and F significantly inhibited the germination of *MoT* conidia. Between the macrolides, oligomycin B showed the lowest conidial germination (24 ± 0.9%) compared to oligomycin F (53±0.4%) at 0.05 μg/ml.

**Table 1.** Effects of oligomycin B and oligomycin F on germination of conidia and their subsequent developmental transitions of *Magnaporthe oryzae Triticum (MoT)* at the dose of 0.05 μg/ml *in vitro*Developmental transitions of the treated conidia of *MoT* by oligomycins

| Compound | Time (h) | Effects of natural macrolides on the developmental transitions of conidia of wheat blast fungus *M. oryzae Triticum* (*MoT*) |  |
| --- | --- | --- | --- |
|  |  | Germinated conidia (% ± SE<sup>a</sup>) | Major morphological change/developmental transitions in the treated conidia |
| Water | 0 | 0 ± 0e | No germination |
|  | 6 | 100 ± 0a | Germinated with normal germ tube and normal appressoria developed |
|  | 12 | 100 ± 0a | Mycelial growth took place |
|  | 24 | 100 ± 0a | Huge mycelial growth took place |
| Oligomycin B | 0 | 0 ± 0e | No germination |
|  | 6 | 38.3 ± 0.7d | 24 ± 0.9% Germinated with a short germ tube and 14.3± 0.7% conidia lysed |
|  | 12 | 24 ± 0.9d | 10.6 ± 0.7% Normal, 8.4 ± 0.2% short and 5 ± 0.4% abnormally elongated germ tube formed |
|  | 24 | 0 ± 0b | No appressoria formed, no mycelial growth took place |
| Oligomycin F | 0 | 0 ± 0e | No germination |
|  | 6 | 59 ± 0.8b | 53 ± 0.4% Germinated with a short germ tube and 6 ± 0.8% conidia lysed |
|  | 12 | 53 ± 0.4b | 33.3 ± 0.5% Normal and 19.7 ± 0.2% abnormal branching at the tips of germ tubes |
|  | 24 | 0 ± 0b | No appressoria formed, no mycelial growth took place |
| Nativo | 0 | 0 ± 0e | No germination |
|  | 6 | 50.3 ± 0.7c | Germinated with a short germ tube |
|  | 12 | 50.3 ± 0.7c | Normal germ tube formed |
|  | 24 | 0 ± 0b | No appressoria formed; no mycelial growth took place |

<sup>a</sup> The data presented here are the mean value ± SE of three replicates in each compound. Means within the column followed by the same letter(s) are not significantly different from those assessed by Tukey's HSD (Honest Significance Difference) post-hoc ( $p \leq 0.05$ ).

### Developmental transitions of the treated conidia of *MoT* by oligomycins

After 24 h of incubation at 25°C in darkness, conidial germination with normal germ tube development and mycelial growth was 100% in water at 6 h, 12 h and 24 h of incubation, respectively (Table 1, Fig 6a). The impact of the two oligomycins on the conidia’s further development differed with time, and abnormal developmental transitions were also found by these natural compounds at 0.05 μg/ml. In case of oligomycin B, 24 ± 0.9% conidial germination took place and 14.3± 0.7% conidia lysis observed after 6 h. At 24 h after treatment, 10.6 ± 0.7% normal, 8.4 ± 0.2% short and 5 ± 0.4% abnormally elongated germ tubes formation observed among the germinated conidia, without progressing appressorial development and mycelial growth after 24 h (Table 1, Fig 6b). In the presence of oligomycin F, 53 ± 0.4% conidia germinated with shorter germ tube and 6 ± 0.8% conidia lysed after 6 h. Among the germinated conidia, 33.3 ± 0.5% normal and 19.7 ± 0.2% abnormally branched germ tube formation took place after 12 h, but no appressoria formation and mycelial growth recorded after 24 h (Table 1, Fig 6c). Nativo induced 50.3 ± 0.7% conidial germination with 50.3 ± 0.7% normal germ tube formation after 6h and 12 h, but inhibited 100% appressoria formation and mycelial growth (Table 1, Fig 6d). The findings revealed that the two macrolides induced major germ tube malformation and suppressed formation of appressoria and hyphal growth of *M. oryzae Triticum.* No conidial malformation was found in case of Nativo^®^WG75, but a slower growth of germ tubes was observed (Fig 6d).

**Fig 6.**
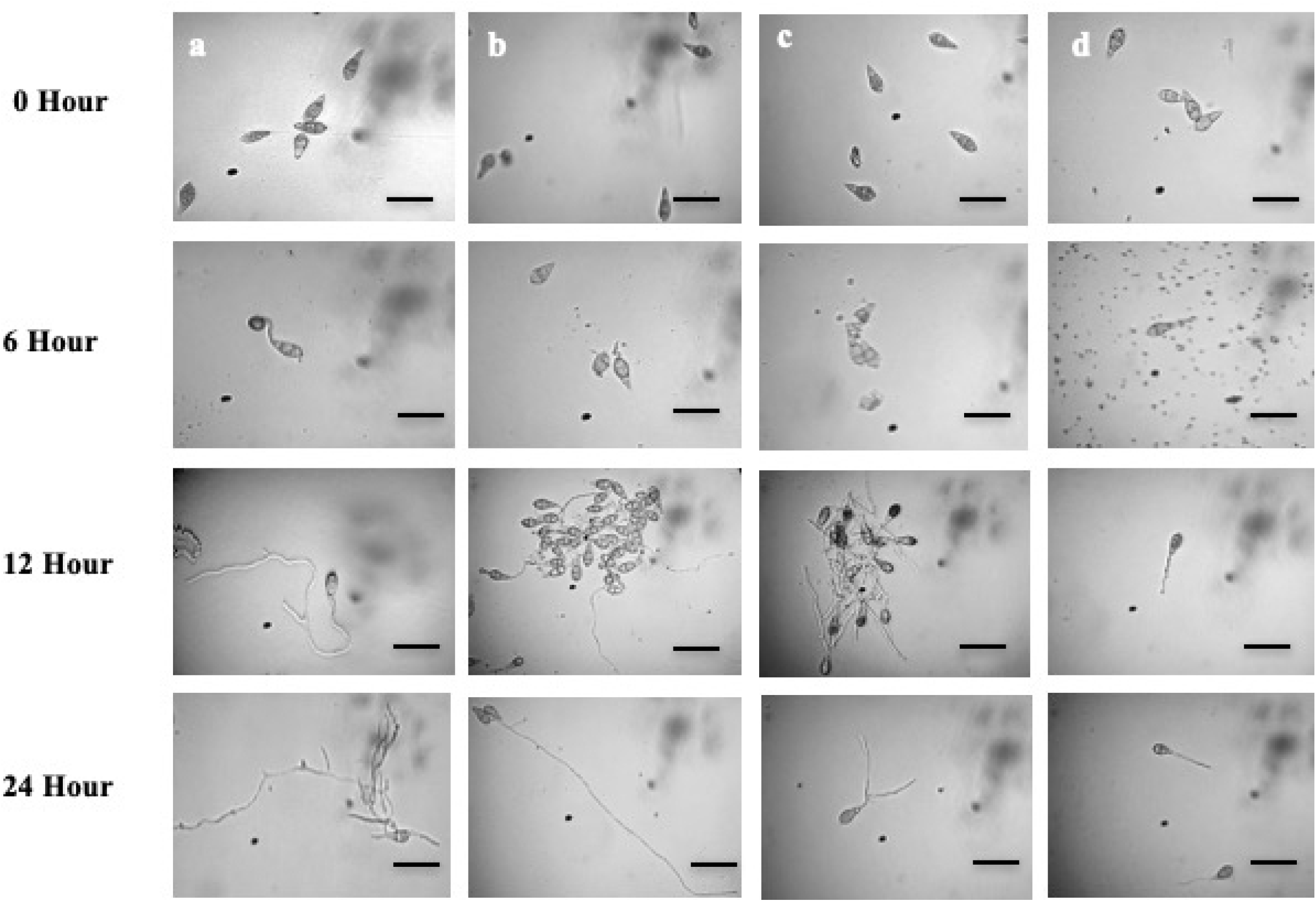
Time-wise alterations in *M. oryzae Triticum (MoT)* conidia germination and their morphological changes in the presence of oligomycin B, oligomycin F and a commercial fungicide Nativo^®^ WG75. Dose of oligomycins was 0.05 μg/ml. (a) Control, (b) Oligomycin B, (c) Oligomycin F, (d) Nativo^®^ WG75. Bar = 10 μm.

### Wheat blast disease suppression by oligomycins

Application of these macrolides at the doses of 5, 10 and 100 μg/ml remarkably inhibited symptoms of wheat blast in detached wheat leaves which were inoculated with *MoT* conidia. The average length of lesions recorded in the wheat leaves treated with oligomycin B were 6.3 ± 0.3 mm and 1.8±0.3 mm at 5μg/ml and 10 μg/ml, respectively (Fig 7A and 7B). In case of oligomycin F and Nativo^®^WG75, recorded blast lesion lengths were 4 ± 0.3 mm and 2 ± 0.3 mm at 5 μg/ml (Fig. 7A, B). No visible disease symptoms were observed on the leaves when treated with oligomycin F and the positive control Nativo^®^WG75 at 10 μg/ml and 100 μg/ml (Fig 7A and 7B). Oligomycin B completely inhibited blast lesion formation at 100 μg/ml. These results show that the development of wheat blast on detached wheat leaves was substantially inhibited by all the natural compounds but at varying doses. On the other side, un-inoculated negative control plants with water treatment had typical blast disease lesions of 9.58 ± 0.2 mm in length (Fig 7A, 7B).

**Fig 7.**
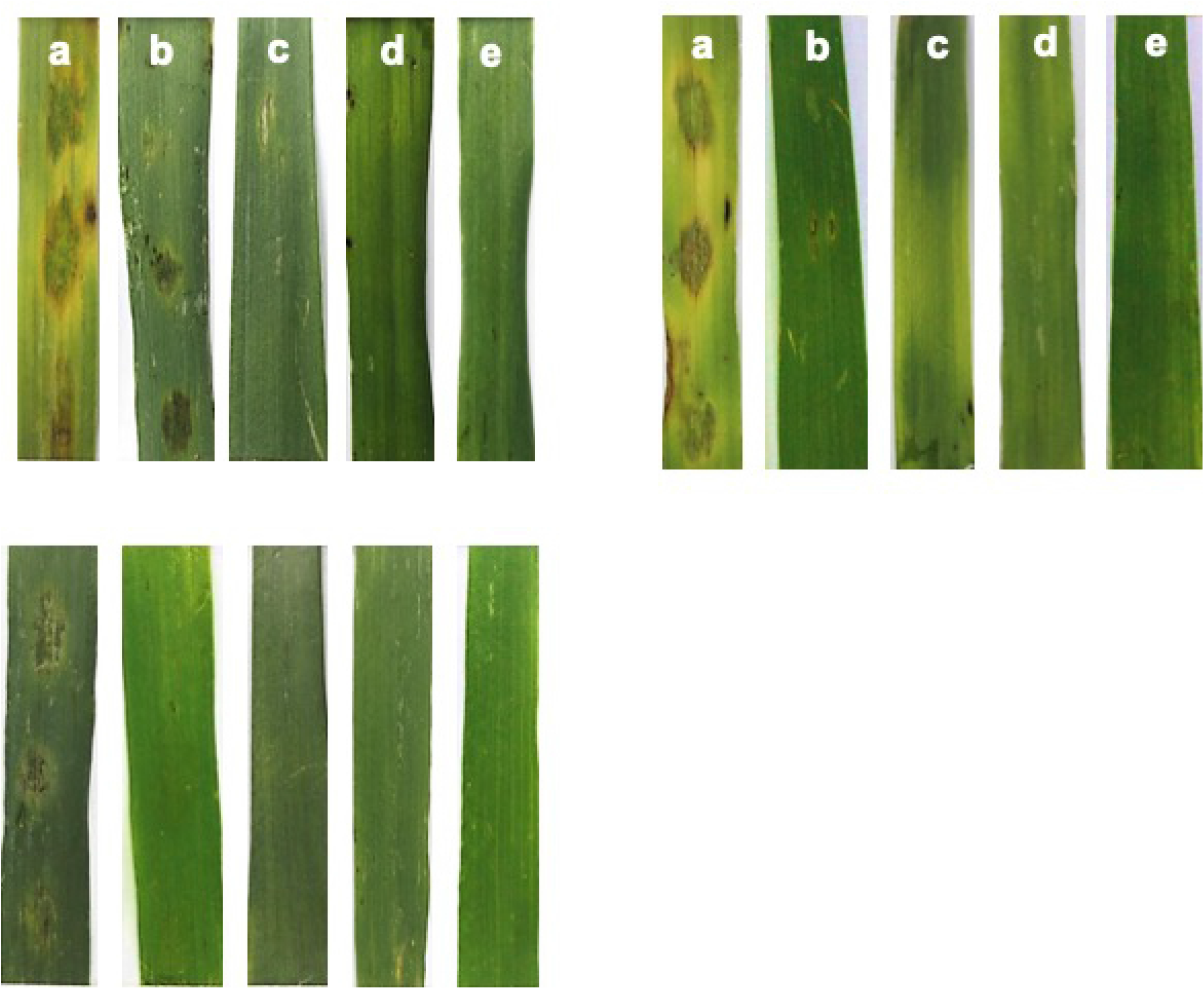
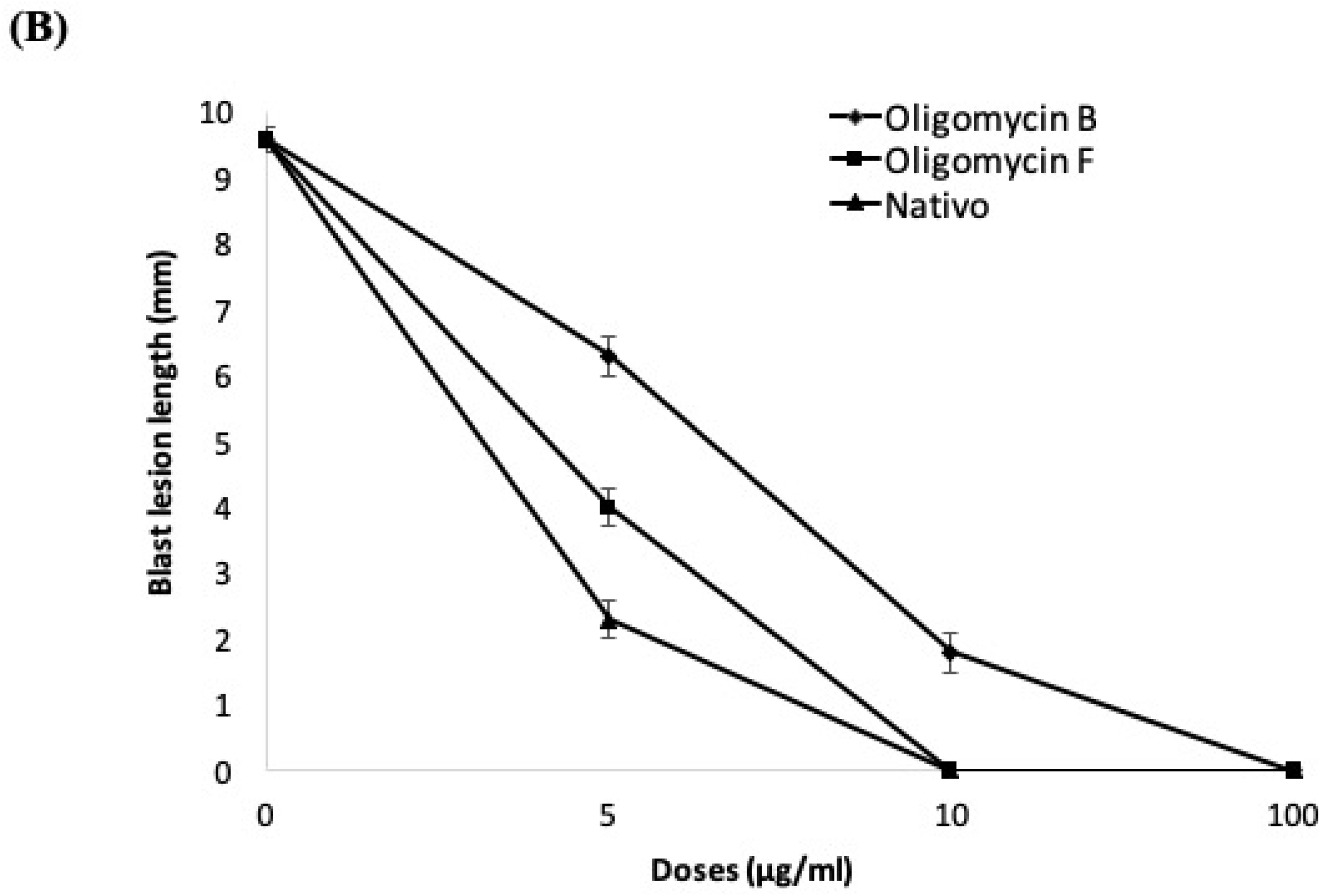
Effects of oligomycins on the suppression of lesion formation in detached wheat leaves by *M. oryzae Triticum* at 5 μg/ml, 10 μg/ml and 100 μg/ml. (A) blast lesion symptoms on treated and untreated wheat leaves (a) Control, (b) Oligomycin B, (c) Oligomycin F, (d) Nativo^®^ WG75, (e) Healthy leaf; (B) diameter of lesions were measured. Bars represent ± standard error. Mean value of control is same for all the chemicals. The data are the averages ± standard errors of at least five replicates for each dose of the tested compounds at p ≤ 0.05.

## Discussion

In this study, we found that two macrolides, oligomycin B and oligomycin F, isolated from *Streptomyces* spp. demonstrated extensive antifungal activities against a devastating blast pathogen of wheat, *M. oryzae Triticum* (*MoT*). Bioassay depicted that both the compounds strongly suppressed mycelial growth as well as conidiogenesis, germination of conidia and morphological changes of the germinated conidia. These natural compounds inhibited wheat blast disease in detached leaves of wheat. The biological activities of the oligomycin F was even stronger than the commercial fungicide Nativo^®^ WG75. Taken together, these findings indicate that suppression of mycelial growth and conidial germination by these macrolides is correlated with wheat blast disease inhibition following inoculation of the leaves. Hyphal growth Inhibition, conidia formation and germination of conidia of various fungi, such as rice blast fungus (*MoO*) by a number of natural secondary metabolites on *in vitro* bioassay have been documented by earlier investigators [3, 11, 34–39]. This is a first report of managing the devastating wheat blast fungus by oligomycin B and F isolated from the *Streptomyces* spp. Oligomycins are macrolide antibiotics that block the proton channel (F0 subunit) requisite for the oxidative phosphorylation of ADP to ATP to inhibit ATP synthase. [40]. While oligomycins have excellent biological properties, only a limited studies have so far focused on their development as plant defense agents. Furthermore, this is also a first report on the inhibition of the wheat killer fungus *MoT* by two macrolide antibiotics, oligomycin B and F from *Streptomyces* spp. Interestingly, the potency of oligomycin F in controlling wheat blast fungus was 10-fold stronger than the commercial fungicide Nativo ^®^ WG75.

One of the striking findings in this present study is the swelling inducing by these macrolides on the hyphae of *MoT* (Fig 2b and 2c). We tested a range of concentrations from 0.005 to 2 μg /disk. Swelling in hyphae increased with increasing concentrations of the oligomycins. Swelling in the fungal hyphae has been reported earlier by polyoxin B [41], fengycin [37–39] and tensin [42]. Morphological changes like extensive branching and swelling of hyphae of an oomycete pathogen *Aphanomyces cochlioides* by phloroglucinols extracted from *Pseudomonas fluorescence* or xanthobaccin A isolated from *Lysobacter* sp. SB-K88 have been documented [43–46]. According to a report of Kim et al. [11], oligomycin A from *Streptomyces libani,* significantly inhibited mycelial growth of *Magnaporthe grisea, Botrytis cinerea, Colletotrichum lagenarium, Cylindrocarpon destructans, Cladosporium cucumerinum and Phytophthora capsici.* So far, this is a first report on swollen-like structures development in hyphae by oligomycins toward the most destructive phytopathogen, *MoT*. A further investigation is needed to understand the mode of action of these macrolides toward the worrisome phytopathogen *MoT*.

Conidia is the medium by which most pathogenic fungi infect plants and the process by which conidia are formed is known as conidiogenesis [35, 47]. Destruction of conidiogenesis and inhibition of conidial germination reduce the chance of fungal phytopathogen infection. Another interesting finding of this study is that both oligomycin B and F strongly suppressed conidiogenesis (Fig 5), germination of conidia and major morphological changes of conidia (Table 1, Fig 6). Bioassay findings showed that previously treated wheat leaves with these macrolides at 5, 10 and 100 μg/ml, resulting in the leaves were less appealing to *MoT* conidia. Others novel phenomena observed in the *MoT* conidia interacting with these macrolides include the dynamics of the lysis of conidia, irregular branching at the tip of germinated conidial germ tube, and abnormally elongated hypha-like germ tube (Fig 6b and 6c). Similar phenomenon was observed by Dame et al. [4] who reported that oligomycin A, B and F from a marine *Streptomyces* can induce lysis of zoospores of a pathogen *Plasmopara viticola* of grapevine downy mildew. Homma et al. [48] reported that, lecithin induces abnormal branching at the conidial germ tube tips and inhibits appressoria formation of rice blast fungus. Islam and Fukushi [46] have observed that cystospores of *A. cochlioides* produced in the presence of diacylphloroglucinol (DAPG) germinated with hyperbranched germ tubes. Carver et al. [49] found that when conidia of *Blumeria graminis* germinated within established oat mildew colonies, most of them developed abnormally elongated hypha-like germ tubes, resulting in no development of appressoria and were unable to attempt invasion. Inhibition of conidiogenesis, germination and formation of appressoria of the *MoT* conidia by the effects of oligomycins has not been reported. These compounds might influence gene expression upstream of melanin synthesis and block intracellular signaling or signal transduction pathways which regulate appressoria formation [50]. The effect of these bioactive natural compounds on the expression of germination of conidia and formation of appressorium related genes in *MoT* will also be focused in future study.

The ATPase has recently appeared as a promising molecular target for developing new therapeutic options for a variety of diseases. The ATP synthase is thought to be one of the oldest and most conserved enzymes in the molecular world and has a complex structure with the probability of inhibition by a range of inhibitors [51]. It is of great scientific concern that oligomycins are macrolide antibiotics that impede ATP production by influencing oxidative phosphorylation in mitochondria [52]. Oligomycin comprises a 26-membered α, β-unsaturated lactone with a conjugated diene fused to a bicyclic spiroketal ring system. Their mode of action includes the decoupling of mitochondrial ATPase F0 and F1 factors responsible for promoting the transfer of proton via the inner mitochondrial membrane [53]. The enzymatic complex F0F1 ATP synthase may be considered as a target for antifungal and anti-tumor or anti-infection therapy [54]. Oligomycins display a number of important biological activities including mitochondrial ATPase inhibition, strong antifungal, anti-actinobacterial and anti-tumor effects have been reported [9, 1, 54]. These natural products are among the strongest selective agents in the cell line; they interrupt P-glycoprotein activity and induce apoptosis in doxorubicin-resistant HepG2 cells [55]. Oligomycins have a variety of isomers called oligomycin A through G. These are particularly relevant to the disruption of mitochondrial metabolism [10]. Moreover, elucidation of structure has provided new horizons for developing new ATP synthase-directed agents with possible therapeutic effects [51]. The first reports of chemical modification of oligomycin A have already been documented by Lysenkova et al. [56]. New compounds have also showed efficacy against *Candida albicans, Aspergillus niger* and *Cryptococcus humicolus*, and their other biological properties are similar to those of oligomycin A, but with less cytotoxic effects. During germination, conidia might need a constant energy (ATP) supply from the internal energy reserve of the cells [57]. It is therefore rational to suppose that the suppression of hyphal growth and conidia germination of *MoT* depicted in this study is likely to be associated with the ATP synthesis inhibition in mitochondria due to the effects of oligomycins. While structural-activity relationships have not been established in the case of these metabolites, more studies are required to determine the precise structure-activity relationships of these oligomycins, which will make it possible to synthesize a more active oligomycin that could be an effective agrochemical against *MoT*.

A hallmark of the findings of this study is that the application of both the macrolides significantly inhibited blast disease development in detached leaves of wheat (Fig 7). In this analysis, the leaves of wheat treated with oligomycin B and F had comparatively shorter lesion lengths than the untreated control (Fig 7). Many of the lesions were small, brown, with pinhead-sized specks (scale 1) to small, roundish to slightly elongated blast lesions infecting <10% of wheat leaf area (scale 5). In contrast, the control wheat leaves had typical blast lesions infecting 26-50% wheat leaf area (scale 7) as per the 9-scale blast disease assessment provided by the IRRI Standard Evaluation System [58]. However, no visible blast lesions were found in the presence of all the compounds and the positive control Nativo ^®^ WG75 at the highest treated concentration (Fig 7). Nativo ^®^ WG75 is a systemic wide-spectrum commercial fungicide that we used as a positive control. Interestingly, the antifungal effect of oligomycins on the inhibition of *MoT* fungus is even equivalent or stronger to that of this fungicide. Tebuconazole and trifloxystrobin are two main active ingredients of Nativo ^®^ WG75. Tebuconazole is known as a dimethylase inhibitor (DMI), which is a systemic triazole fungicide. This interacts with sterol biosynthesis in fungal cell walls and suppresses spore germination and further development of the fungus [59]. Trifloxystrobin is a strobilurin fungicide that interferes with the respiration of plant pathogenic fungi by preventing electron transmission in mitochondria, disrupting metabolic processes and inhibiting the development of target fungi [60]. The mechanism of disease suppression by the oligomycins might be uniquely different from the Nativo ^®^ WG75. A further study is needed to elucidate the underlying mechanism of wheat blast disease suppression by the oligomycin B and F. Furthermore, a field trial of the oligomycins in controlling wheat head infection is needed before considering them as effective fungicides against the wheat blast.

Despite the significant potency of an antifungal antibiotic, little has been reported about the effectiveness of oligomycins as agricultural fungicides. The shorter residual effect of oligomycin can be important in the reduction of deleterious effects on humans and the environment, considering that sufficient efficacy in the management of plant diseases is sustained [11]. In the *in vivo* experiments for the protection efficacy against plant diseases such as *Phytophthora* blight in pepper plants, anthracnose in cucumber leaves, and leaf blast in rice, oligomycin A demonstrated substantially defensive activity against certain plant pathogen infections in host plants treated with 500 μg/mL. Defense activity at this concentration was extended for a month, resulting in no signs of disease on host plants [11]. Oligomycin A was found to be the most active anti-filamentous fungi analogue among antibiotics in the oligomycin family [1, 61, 62]. Since oligomycin F is the immunosuppressive homolog of oligomycin A [3] and oligomycin B is a stable natural product [63], these macrolides might be the leading compounds for the production of agrochemicals against the cereal killer *MoT*.

The morphological changes seen by these macrolides in *MoT* are also linked to several active, selective and cell-permeable antifungal secondary metabolites, like chelerythrine chloride, staurosporin, polyoxin A, polyoxin B and polyoxin D. Chelerythrine chloride and staurosporin are inhibitors of protein kinase C that have induced swelling of hyphal subapical areas, inhibited spore germination, and inhibited the formation of appressoria of plant pathogenic fungi by blocking ATP binding to kinase [64–67]. Polyoxins are well-known as inhibitors of cell wall biosynthesis that suppress the formation of germ tubes and appressoria [41, 68–70]. The findings of this research were also comparable to the activities of several commercially available fungicides such as blasticidin-S, kasugamycin, streptomycin and tricyclazole, which demonstrated strong antifungal efficacy against many phytopathogenic fungi, like rice blast fungus *M. oryzae Oryzae (MoO)* [11, 71–73]. This study for the first time showed that two secondary metabolites of *Streptomyces* spp. were used to suppress the wheat blast disease. The typical mode of action of these fungicides are the inhibition of protein synthesis, microcondrial ATPase, cell wall biosynthesis and melanin biosynthesis, resulting in the inhibition of fungal mycelial growth, germination of spore and formation of appressoria.

Nowadays, the increase in resistance to fungicides among pathogenic micro-organisms is an alarming problem in agriculture. Non-rational application of commercial fungicides with sitespecific modes of action, like strobilurin (QoI) and triazole, has resulted in the widespread dissemination of several resistant mutant species in *MoT* [27, 28]. The most concerning antifungal resistance problem of conventional fungicides contributes to a search for new, effective antifungal agents to protect wheat plants against this dangerous phytopathogenic fungus. The macrolides, oligomycin B and F have a comparatively higher bioactivity compared to commercial fungicide Nativo ^®^ WG75. The inhibitory ability of these macrolides will allow them to be considered as candidate agrochemicals with a novel mode of action against the notorious wheat blast fungus.

## Conclusion

Our findings for the first time, have shown that the two macrolides, oligomycin B and F from *Streptomyces* spp., suppressed hyphal growth and asexual development of *MoT* fungus, and inhibited wheat blast disease development in detached leaves of wheat. Field assessment of these macrolides is required to evaluate these metabolites as effective fungicides against wheat blast disease. Further research is also required to understand the mode of action and the structureactivity relations among oligomycins A-G towards a devastating wheat killer, *M. oryzae Triticum*.

## Funding

This work was funded by the Krishi Gobeshona Foundation (KGF), Bangladesh through a coordinated project No. KGF TF 50-C/17 to Tofazzal Islam of the Institute of Biotechnology and Genetic Engineering of BSMRAU, Bangladesh.

## Conflict of Interest Statement

The authors declare that there is no conflict of interest.

## Ethics statement

There is no ethical issues in the experiment of this manuscript.

## Author contributions

**Conceptualization:** Tofazzal Islam

**Data curation:** Tofazzal Islam

**Formal analysis:** Moutoshi Chakraborty, Nur Uddin Mahmud

**Investigation:** Moutoshi Chakraborty, Nur Uddin Mahmud

**Resources:** Moutoshi Chakraborty, Nur Uddin Mahmud, Abu Naim Md. Muzahid, S. M. Fajle Rabby

**Methodology:** Moutoshi Chakraborty, Nur Uddin Mahmud, Abu Naim Md. Muzahid, S. M. Fajle Rabby

**Project administration:** Tofazzal Islam

**Software:** Moutoshi Chakraborty, Nur Uddin Mahmud

**Supervision:** Tofazzal Islam

**Funding acquisition:** Tofazzal Islam

**Visualization:** Tofazzal Islam

**Validation:** Tofazzal Islam.

**Writing - original draft:** Moutoshi Chakraborty, Nur Uddin Mahmud

**Writing - review & editing:** Tofazzal Islam

## Acknowledgements

The authors are thankful to the Krishi Gobeshona Foundation (KGF), Bangladesh (KGF-TF 50C/17) for funding this work. The funders had no role in study design, data collection and analysis, decision to publish, or preparation of the manuscript. The authors are also thankful to Dr. Hartmut Laatsch of Georg-August University Goettingen, Germany, for kindly providing the oligomycins for this research.

